# In utero delivery of miRNA induces epigenetic alterations and corrects pulmonary pathology in congenital diaphragmatic hernia

**DOI:** 10.1101/2022.02.27.482144

**Authors:** Sarah J Ullrich, Nicholas K Yung, Tory J Bauer-Pisani, Nathan L Maassel, Mary Elizabeth Guerra, Mollie Freedman-Weiss, Samantha L Ahle, Adele S Ricciardi, Maor Sauler, W Mark Saltzman, Alexandra S Piotrowski-Daspit, David H. Stitelman

## Abstract

Structural fetal diseases, such as congenital diaphragmatic hernia (CDH) can be diagnosed prenatally. Neonates with CDH are healthy *in utero* as gas exchange is managed by the placenta, but impaired lung function results in critical illness from the time a baby takes its first breath. During fetal development, lungs are capable of remarkable growth and the fetus does not yet require lung function for gas exchange. MicroRNA (miR) 200b and its downstream targets in the TGFβ pathway are critically involved lung branching morphogenesis. Here we characterize the expression of miR200b and the TGFβ pathway at different gestational times using a rat model of CDH. Fetal rats with CDH are deficient in miR200b at gestational day 18. We demonstrate that NPs loaded with miR200b given systemically to fetal rats result in changes in the TGFβ pathway; these epigenetic changes improve lung size, lung morphology, and lung vascularization. This is the first demonstration of *in utero* epigenetic therapy to improve lung growth and development in a pre-clinical model. With refinement, this technique could be applied to fetal cases of CDH or other forms of impaired lung development in a minimally invasive fashion.

**eTOC Synopsis:** *In utero* treatment with NPs loaded with miR200b improves lung development in a rat model of CDH. miR200b treatment epigenetic changes in the TGFβ, leads to larger lungs with more airspace and favorable pulmonary vascular remodeling.

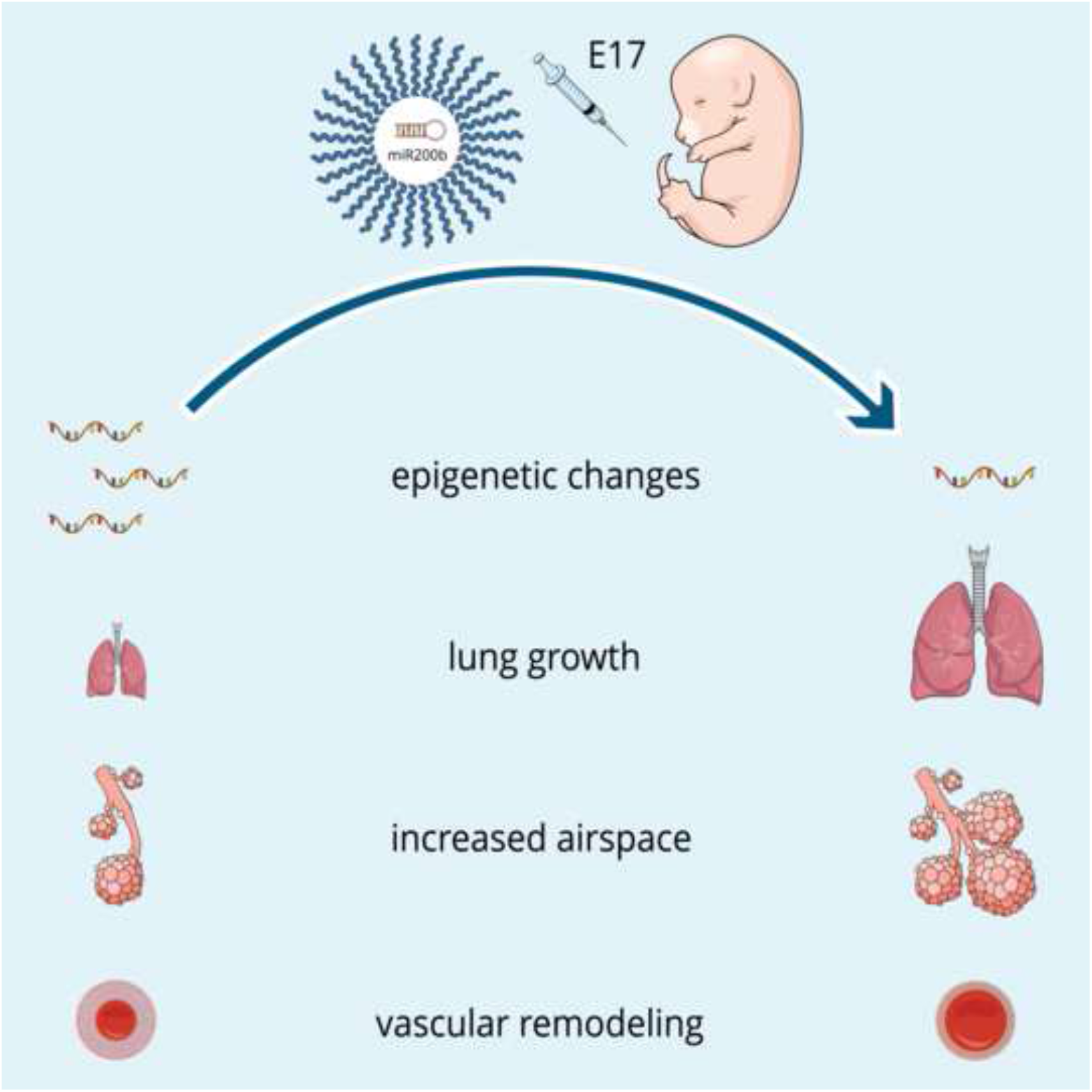

## Introduction

Congenital Diaphragmatic Hernia (CDH) results from incomplete fusion of the diaphragm *in utero* causing the abdominal organs to herniate into the chest cavity.^1,2^ It is a relatively common condition with an incidence of 1 in 2,000 to 3,000 live births.^3,4^ Even with modern neonatal critical care, the overall mortality is approximately 30%, and approaches 80% for those with severe disease.^5,6^ Infants with CDH have poorly developed lungs with two predominant pathologic features: pulmonary hypoplasia (PH) characterized by reduced distal airway branching with fewer alveoli that are thick walled and persistent pulmonary hypertension (PPH) due to hypermuscularized arterioles that are less vasoactive.^7–10^

There is a compelling rationale to develop fetal intervention for CDH given the potential to reverse pulmonary hypoplasia *in utero*, before the delivered fetus takes its first breath. Trials examining Fetoscopic Endoluminal Tracheal Occlusion (FETO), where a balloon is deployed in the trachea under fetoscopic guidance and subsequently removed weeks later, have demonstrated a survival benefit for those with severe disease.^11,12^ But FETO carries a substantial risk of complications such as preterm premature rupture of the membranes and preterm delivery. Among fetuses treated with FETO, those who had subsequent lung growth and survived after balloon deployment and retrieval had higher miR200b expression in their tracheal fluid aspirates than those who did not respond to tracheal occlusion.^13^ The microRNA 200 family inhibits several genes in the TGFβ pathway, which has been previously demonstrated to modulate lung branching during development, and *in vitro* studies suggest that treatment with miR200b can increase branching morphogenesis.^13–18^

miRNAs are rapidly degraded by serum enzymes and intracellular RNases, making them highly unstable *in vivo*.^19^ For miRNAs to be used therapeutically, it is essential for them to be delivered in a vehicle that confers high stability and avoids potential toxicities.^20,21^ A variety of approaches have been explored to address these limitations, such as chemically modifying the miRNA mimics to enhance stability or using viral vectors, lipid emulsions, liposomes, lipid nanoparticles, or polymeric nanoparticles.^19,22–29^ But stabilized miRNA mimics have decreased mRNA silencing ability, viral vectors have a poor safety profile, and cationic lipids are toxic to cells and pro-inflammatory.^19,30^ Polymeric nanoparticles (NPs) can be used for the sustained delivery of drugs, are non-toxic with a low side effect profile, and can be optimized to target fetal lung.^31–33^ We have recently demonstrated that polymeric NPs can safely be used to deliver editing reagents in the form of peptide nucleic acids (PNAs) and donor DNAs *in utero* to correct a disease-causing mutation in the β-globin gene in a mouse model of human β-thalassemia.^34^ Optimization for delivery to fetal lung demonstrated that intravenous delivery led to improved lung delivery over intraamniotic delivery, and that cationic poly(amine-co-ester) (PACE) nanoparticles (NPs) were delivered most efficiently to fetal lung.^33^

Here, we sought to deliver miR200b *in utero* using PACE NPs to treat a rat model of CDH. We found that *in utero* delivery of miR200b induces alterations in the TGFβ pathway, leads to pulmonary vascular remodeling, and improves pulmonary hypoplasia.

## Results

### miR200b expression and TGFβb signaling is altered during late gestation in the Nitrofen model of CDH

Branching morphogenesis of fetal rat lung is completed during the canalicular stage of lung development (E18 through E20), whereas the terminal airspaces develop and divide during the saccular stage (E21-P4).^35^ It has been previously demonstrated that miR200b regulates distal airway branching, and that at E18 miR200b expression is reduced in nitrofen exposed lungs, and at E21 miR200b expression is increased in nitrofen exposed lungs.^17,18^ To determine an ideal therapeutic window for *in utero* miR delivery, we further elucidated the temporal expression of miR200b in the wild-type (WT) and nitrofen-induced CDH lungs. The relative expression of miR200b in WT and CDH lungs were quantified using RT-qPCR. In WT fetal rat lung, miR200b levels are stable during the canalicular phase and undergo a 1.7-fold increase at the transition to the saccular phase (Figure 1A). In CDH lungs, there is a 40% decrease in the relative expression of miR200b at E18 during the early canalicular stage (Figure 1B). During the mid and late canalicular phase, there is no difference in the relative expression of miR200b between WT and CDH lungs. At E21, the relative expression of miR200b is 1.4-fold greater in CDH lungs compared with WT lungs.

**Figure 1:**
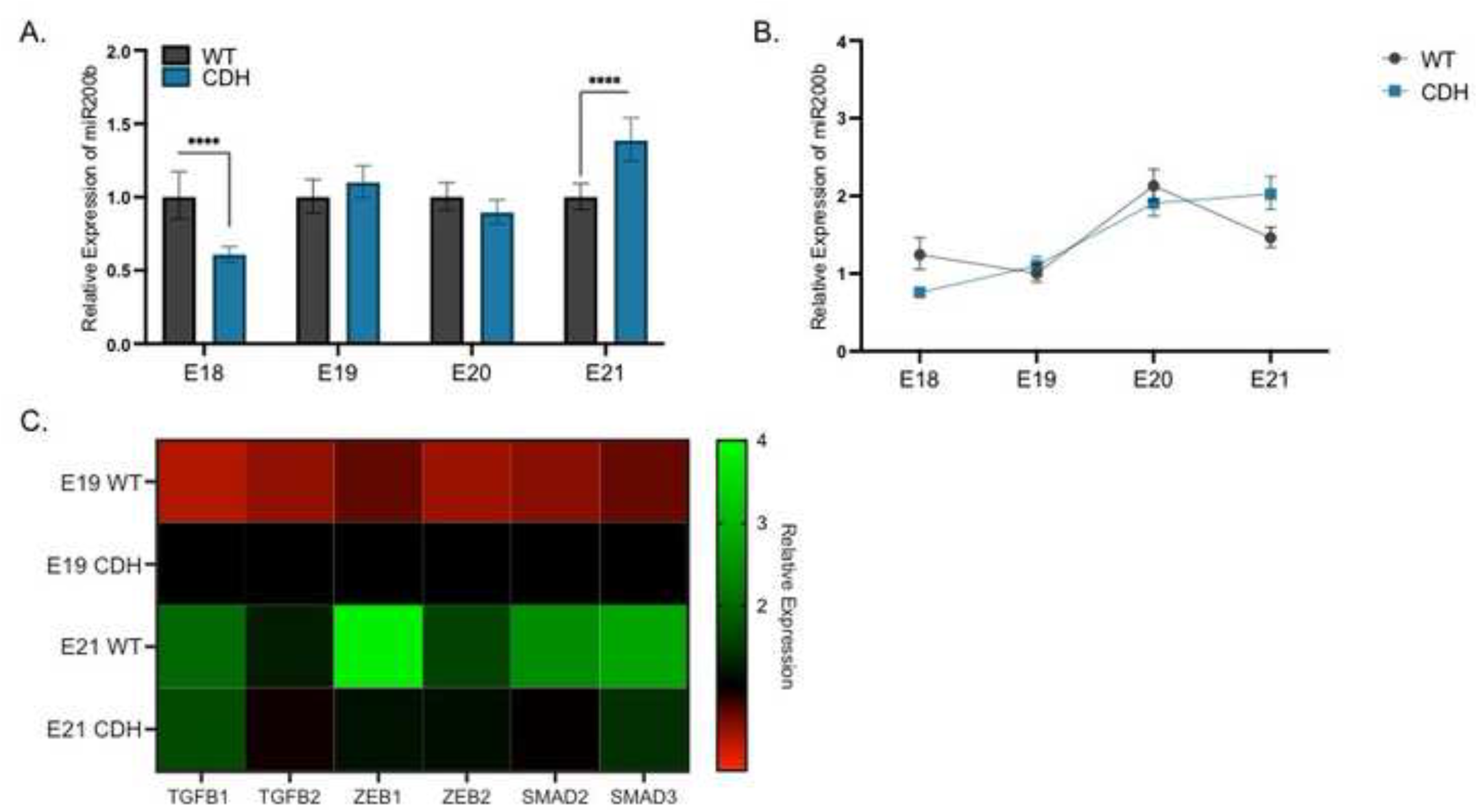
miR200b and the TGFβ pathway expression during the canalicular and saccular phases of lung development in the Nitrofen rat model of CDH. A) Bar graph showing the relative abundance of miR200b in wild type and CDH lungs during late gestation. All values normalized to wild-type at each day. Students t-test,**** p< 0.0001, Data are shown as mean + S.D. of n = 10-12. B) Relative expression of miR200b during late gestation in wild type and Nitrofen induced CDH fetal rat lungs during the canalicular and saccular phases of pulmonary development normalized to wild-type E19 lungs. Data shown as mean + S.D. of n = 10-12 C) Heatmap of the relative expression of the TGFβ pathway during the canalicular (E19) and saccular (E21) phases of pulmonary development in WT and CDH lungs. Red = downregulated, green = upregulated. n=4.

The TGFβ signaling pathway is a regulator of branching morphogenesis in pulmonary development and is modulated by miR200b.^36,37^ The role of the TGFβ pathway in CDH is less clearly defined.^38^ We found that in WT lungs, the expression of TGFβ and its downstream targets increases as the lungs transition from the canalicular to the saccular phase (Figure 1C). In CDH lungs, the expression of TGFβ and its downstream targets is increased during the canalicular phase compared with WT lungs. In CDH lungs, the TGFβ pathway does not undergo the same increase in expression during late development, with the relative expression of most downstream targets remaining stable between E19 and E21.

### In utero delivery of miR200b induces epigenetic changes

Because we found that miR200b levels were reduced in CDH lungs at E18, we elected to target this time point for treatment. Our previous data demonstrated that the majority of nucleic acid release from PACE60 NPs occurs within the first 24 hours.^39^ To align delivery of miR200b with the appropriate therapeutic window, we injected NPs at E17. To validate if *in utero* injection of miRNA PACE60 NPs led to an increase in miR200b levels, WT fetuses were injected with control or miR200b loaded NPs. Lungs were evaluated 4 hours after injection. Those injected with miR200b had a 12 fold increase in miR200b levels (Figure 2A).

**Figure 2:**
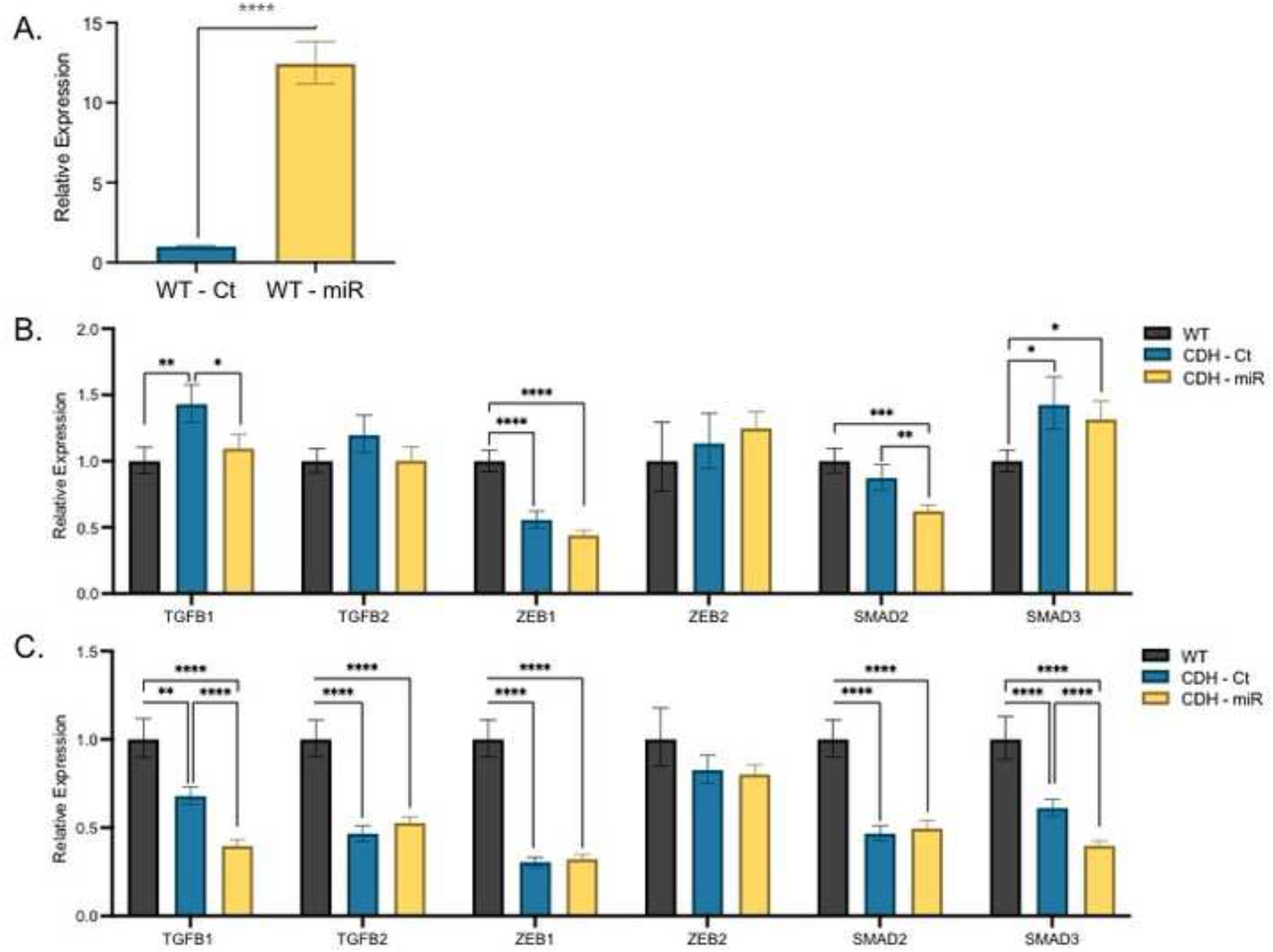
*In utero* delivery of miR200b by PACE60 NPs alters miR200b expression levels and induces epigenetic changes in the TGFβ pathway. A) Bar graph showing the relative abundance of miR200b in WT fetal lungs at E17 4 hours post injection with control and miR200b PACE60 NPs. Values normalized to lungs injected with control miRNA PACE NPs. Students t-test,**** p< 0.0001, Data are shown as mean + S.D. of n = 3. B) Bar graph showing the relative abundance of the downstream targets of the TGFβ pathway in WT fetal lungs and those injected with control miRNA and miR200b PACE60 NPs at E19. Values normalized to WT lungs. 2-Way ANOVA, * p< 0.05, **p<0.01, *** p< 0.001, **** p< 0.0001, Data are shown as mean + S.D. of n = 4-6. C) Bar graph showing the relative abundance of the downstream targets of the TGFβ pathway in WT fetal lungs and those injected with control miRNA and miR200b PACE60 NPs at E21. Values normalized to WT lungs. 2-Way ANOVA, **p<0.01,**** p< 0.0001, Data are shown as mean + S.D. of n = 4-6.

Previous *in vitro* studies demonstrated that treatment with a miR200b mimic abrogates nitrofen induced upregulation of the TGFβ pathway. We treated nitrofen-exposed pups with control or miR200b NPs at E17 and assessed the relative expression of the TGFβ pathway using RT-qPCR at E19 and E21. We found that pups with CDH who were treated with miR200b PACE60 NPs had a decrease in the relative expression of TGFβ1 and SMAD2 at E19 compared with pups treated with control NPs (Figure 2B). At E21, pups treated with miR200b had a decrease in TGFβ1 expression and SMAD3 expression compared to those treated with control NPs (Figure 2C). There were no instances where pups treated with control NPs had a change in gene expression and miR200b PACE60 NPs did not.

### Lung morphology improves after in utero treatment with miR200b

To assess if *in utero* treatment with miR200b lead to morphometric changes in fetal lung, lungs of CDH pups were harvested at E21 after treatment with either control or miR200b PACE60 NPs at E17. Treatment with miR200b improved pulmonary hypoplasia in pups with CDH (Figure 3 A-C). The lung to body weight ratios of pups treated with miR200b NPs increased by 27% (2.2% vs 2.8%), and approached the ratios of WT pups (3.4%, Figure 3D). Pulmonary airspace improved 140% in pups treated with miR200b, approaching that of WT pups, as measured by mean linear intercept (WT 37μM, control 11μM, miR200b 26μM, Figure 3E). Additionally, pups treated with miR200b had a 63% decrease in mean alveolar thickness, decreased beyond that of wild type pups (WT 9μM, control 16μM, miR200b 6μM, Figure 3F).

**Figure 3:**
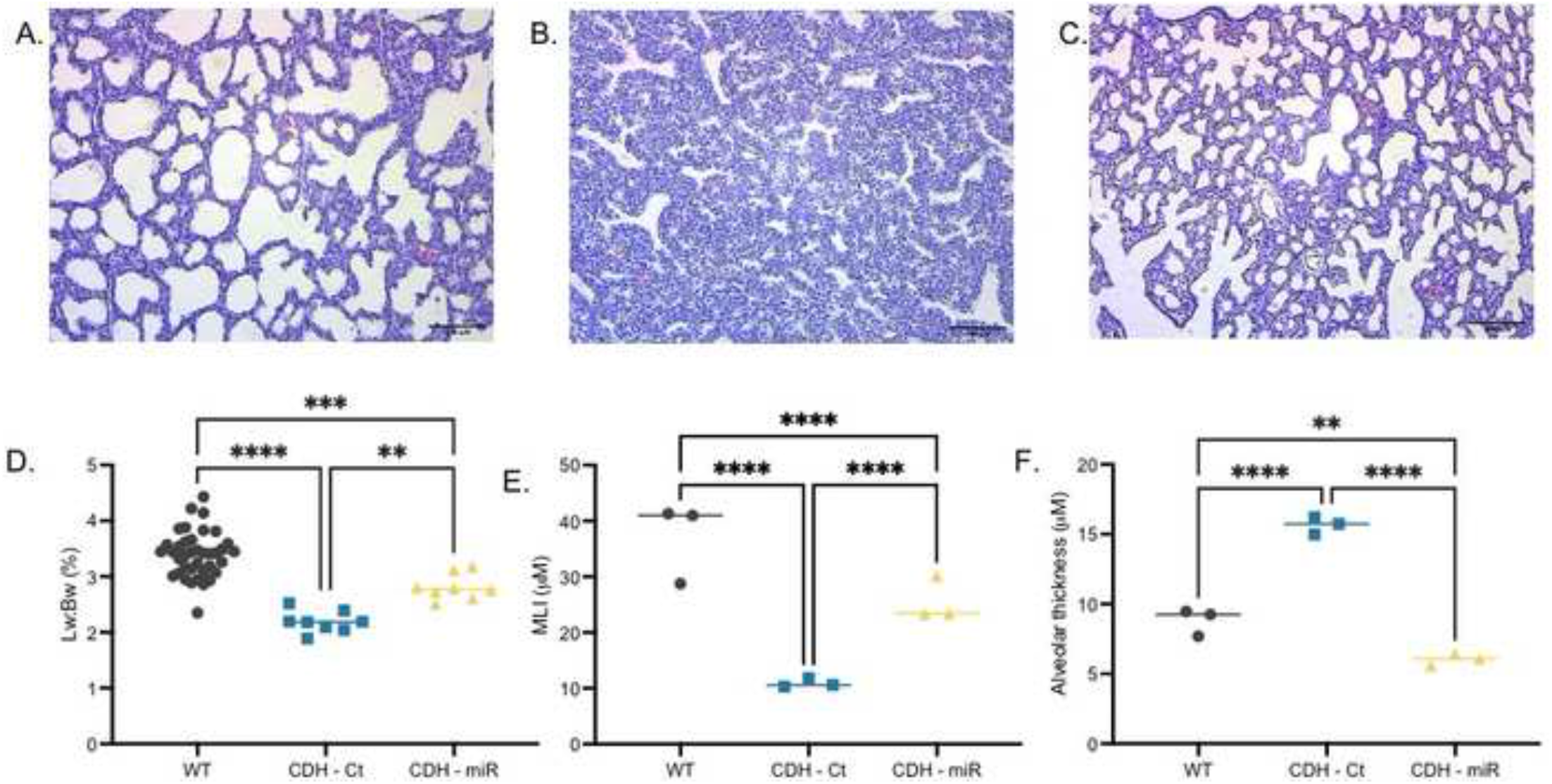
*In utero* delivery of miR200b PACE60 NPs improves pulmonary hypoplasia in the Nitrofen Rat model of CDH Representative hematoxylin and eosin images of A) Term WT lungs. B) Term CDH lungs treated with Control miRNA PACE60 NPs and C) Term CDH lungs treated with miR200b PACE60 NPs. Scale bars = 100 μm. D) Lung to body weight ratios of WT lungs, CDH lungs treated with control miRNA PACE60 NPs and CDH lungs treated with miR200b PACE60 NPs. Each data point represents a single pup. 1-Way ANOVA, **p<0.01, *** p< 0.001, **** p< 0.0001. E) Mean linear intercept of WT lungs, CDH lungs treated with control miRNA PACE60 NPs and CDH lungs treated with miR200b PACE60 NPs. Each data point represents the lung of a single pup. 1-Way ANOVA, **** p< 0.0001. F) Mean alveolar thickness of WT lungs, CDH lungs treated with control miRNA PACE60 NPs and CDH lungs treated with miR200b PACE60 NPs. 1-Way ANOVA, **p<0.01, *** p< 0.001, **** p< 0.0001.

*In utero* treatment with miR200b also led to pulmonary vascular remodeling (Figure 4A-C). The pulmonary arteries of pups injected with miR200b had a 23% decrease in percent medial wall thicknesses compared with those treated with control NPs (37.3% vs 48.5%, Figure 4D). The pulmonary arteries of pups treated with miR200b showed equivalent percent medial thicknesses to WT pups (Figure 4D).

**Figure 4:**
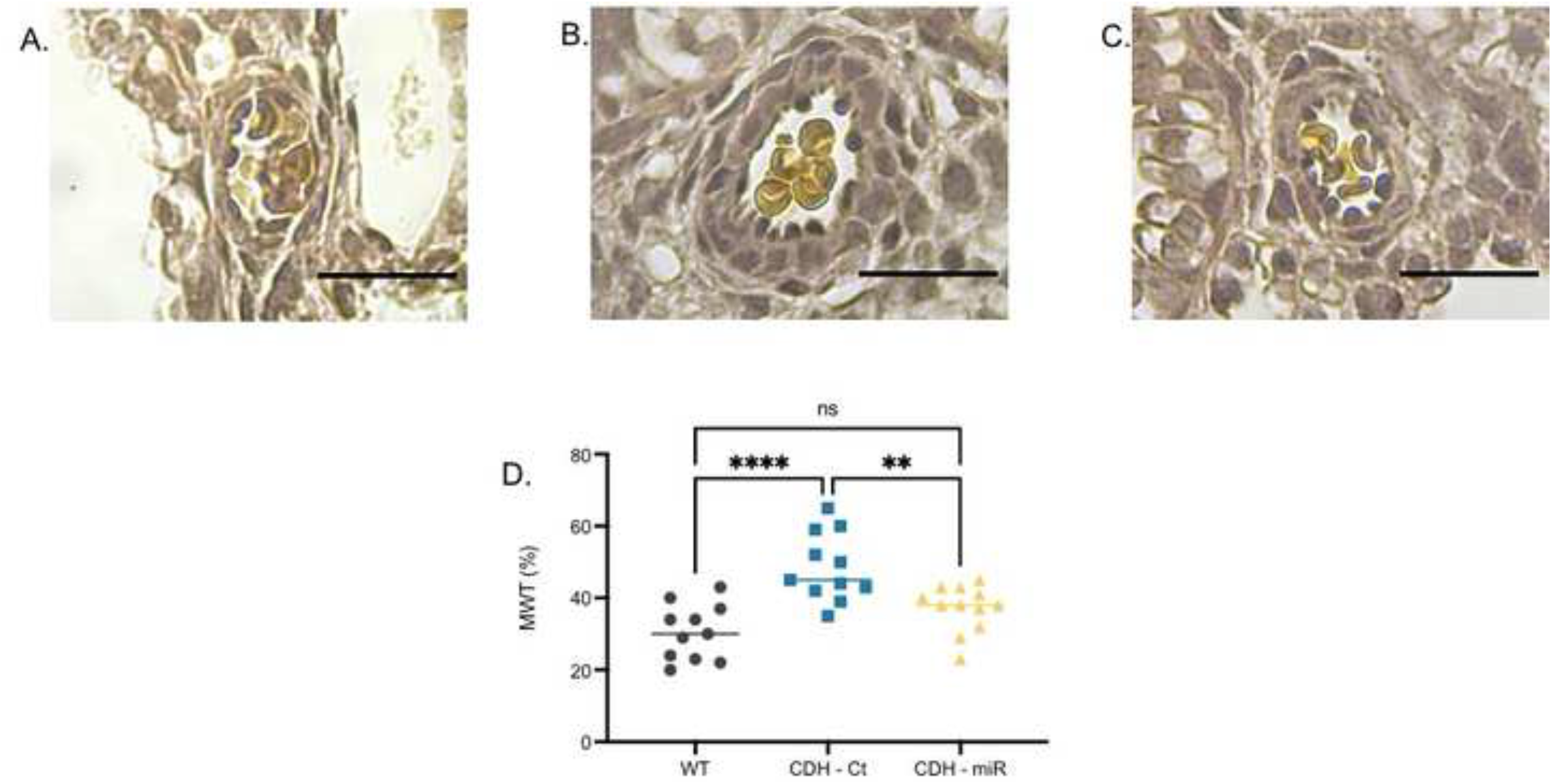
*In utero* delivery of miR200b PACE60 NPs results in pulmonary vascular remodeling in the Nitrofen Rat model of CDH. Representative images of Elastin van Gieson stained pulmonary arteries from A) Term WT lungs B) Term CDH lungs treated with control miRNA PACE60 NPs and C) Term CDH lungs treated with miR200b PACE60 NPs. Scale bars = 20 μm. D) Percent medial wall thickness of WT lungs, CDH lungs treated with control miRNA PACE60 NPs and CDH lungs treated with miR200b PACE60 NPs. Each data point represents an individual arteriole. 1-Way ANOVA, **p<0.01, **** p< 0.0001.

## Discussion

In this study, we demonstrated that a single intravenous fetal dose of PACE60 NPs loaded with miR200b improves lung development in a rat model of CDH. Specifically, delivery of miR200b induced epigenetic changes in the TGFβ pathway during the canalicular and saccular stages of pulmonary development. *In utero* treatment with miR200b lead to larger lungs with more airspace and favorable pulmonary vascular remodeling at term in pups with CDH. This targeted, minimally invasive *in utero* strategy has the potential to avoid substantial antenatal morbidity and mortality in CDH.

We examined the relative expression of miR200b during the canalicular and saccular stages of rat lung development in both WT and CDH lungs and found that there is a marked deficiency in miR200b at E18, followed by a rise in relative expression leading to overexpression at E21. The relative expression of miR200b was equivalent at E19 and E20, suggesting the early canalicular phase is an optimal time for delivery of miR200b. CDH is a devastating lung disease defined by impaired pulmonary vasculature and distal airway branching morphogenesis. miR200b plays a key role in this process, promoting distal airway development by maintaining an epithelial cell phenotype.^17,18,36,40^ The role of miR200b in the pathogenesis of CDH has been less clearly delineated. Here, we defined a key role for miR200b in the pathogenesis of CDH. Our findings are consistent with previous findings which examined miR200b expression in nitrofen exposed lungs with varying degrees of hypoplasia.^18^

We demonstrated that in the nitrofen rat model, the TGFβ pathway is upregulated in the lungs of pups with CDH relative to WT pups during the canalicular stage of lung development. During the transition from the canalicular to saccular stages, the TGFβ pathway is upregulated as branching morphogenesis slows down. However, in pups with CDH, the expression of the TGFβ pathway does not change significantly, further indicating that the dynamics of the TGFβ pathway is dysfunctional in the lungs of pups with CDH. The TGFβ pathway is one of many signal transduction pathways regulating branching morphogenesis in the developing lung. TGFβ signaling is particularly important at the epithelial-mesenchymal interfaces during fetal development and plays a key role in branching morphogenesis.^16^

In this study, we demonstrated that *in utero* treatment with miR200b can induce epigenetic changes in the fetal lung. Though treatments designed to modify tumor epigenetics have long been investigated for cancer, this has not been a focus of fetal interventions.^41^ Treatment strategies that induce fetal epigenetic changes are particularly promising for CDH where – unlike other congenital pulmonary diseases such as cystic fibrosis—a monogenic cause has not been identified.^42^ This suggests that environmental, epigenetic factors play a substantial role in the pathophysiology of CDH.

Fetal therapy for CDH, before a baby takes a first breath, is the optimal time to improve lung development as the lung is capable of structural change during fetal life and the gas exchange is managed by the placenta. We envision that NP-based fetal epigenetic therapy could be delivered through a small needle via ultrasound guided cannulation of the umbilical vessels, which has been in clinical practice since the 1980’s and carries a low risk of fetal loss (~1%).^43–45^ Though FETO has been demonstrated to have a survival benefit for those with severe disease, it is an invasive procedure, and is associated with substantial risk of complications such as preterm premature rupture of the membranes and preterm delivery. Another major limitation of FETO is that while it induces lung growth, it does not improve pulmonary vascular remodeling and pulmonary hypertension.^46^ In this study, we demonstrated that *in utero* treatment with miR200b PACE60 NPs leads to favorable pulmonary vascular remodeling, in addition to improving pulmonary hypoplasia.

Though the PACE NPs used for this study have been optimized for delivery to fetal lung, there is still the potential for off-target effects due to delivery to off target tissues.^33^ As this therapy is translated to larger animal models, direct intra-tracheal administration could be considered to minimize off-target delivery; intratracheal PACE vehicle delivery is well tolerated in adult animals.^47^ We have demonstrated that miR200b delivery induces epigenetic changes in the TGFβ pathway, but it also impacts other signal transduction pathways; further study is warranted before translation to larger animal models.^15^ Success in such investigations could provide a strong foundation for clinical translation of this novel therapeutic modality.

## Materials and Methods

### Polymer Synthesis and Nanoparticle Formulation

PACE polymers were synthesized as described previously.^39^ Briefly, monomers (15-pentadecanolide, N-methyl diethanolamine, and diethyl sebacate) were dissolved in diphenyl ether with an enzyme lipase catalyst (Novozym 435). 60 mol% PDL was used for the PACE polymer generated for this study. The synthesis reaction proceeded in two phases: an oligomerization phase for 18-20 hours under argon at 1 atm, and then a polymerization phase for 48-72 hours under vacuum. PACE polymers were characterized by NMR and GPC. PACE60 nanoparticles were formulated as described previously by double emulsion solvent evaporation using miR200b mimic (Invitrogen *mirVana* miRNA Mimic, hsa-miR-200b-3p, 5’-UAAUACUGCCUGGUAAUGAUGAC −3’, Cat# 4464066, Ambion, Austin, TX) or a negative control miR (Invitrogen *mir*Vana miRNA Mimic, Negative Control #1, Cat# 4464059, Ambion, Austin, TX).^39,48^ Characteristics of PACE60 NPs are displayed in Table 1.

**Table 1.** Characterization Data for PACE60 NP formulations.

| Formulation | Diameter (nm) | Polydispersity Index (PDI) | Zeta Potential (mV) |
| --- | --- | --- | --- |
| PACE60 control miRNA | $301 \pm 3$ | 0.170 | $30.2 \pm 0.1$ |
| PACE60 miR200b | $323 \pm 2$ | 0.190 | $31.2 \pm 0.2$ |

### Rat Model and Injection

All animal use was in accordance with the guidelines of the Animal Care and Use Committee of Yale University. CDH was modeled in rats as previously described.^49^ Time-dated pregnant Sprague–Dawley rats from Charles River Laboratories (Wilmington, MA) were gavage fed 100 mg of Nitrofen (Cat# 33374, Sigma Aldrich, St. Louis, MO) dissolved in olive oil (Whole Foods, Austin, TX) on embryonic day E9. Time dated pregnant rats at 17 days post conception were anesthetized with inhaled isoflurane (2% vol/vol for induction and maintenance). Analgesia was provided with subcutaneous injection of 0.3mL Buprenorphine HCl (0.015mg/mL) and 0.3mL Meloxicam (0.1mg/mL). The gravid uterus was exposed through a midline laparotomy incision. miR-loaded PACE60 NPs were resuspended by vortex and water bath sonication and delivered a dose of 30 mg/kg in 1× DPBS. NP suspension was drawn up into a glass micropipette (tip diameter ~60 μm) and injected intravascularly via vitelline vein of each fetus using a pneumatic microinjector (Narishige; Japan). Rats were sacrificed at varying time points post injection and fetuses were delivered by cesarean section.

### RT-qPCR

Total miRNA from fetal rat lung was extracted using the mirVana miRNA Isolation Kit (Cat# AM1560, Ambion, Austin, TX, USA) according to the manufacturer’s instructions. RT-qPCR was performed for miRNA 200b and its mRNA downstream targets.

For the miRNA assay, 1-10 ng total RNA were reverse transcribed using the TaqMan miRNA Assay for rno-miR200b-3p (rno481286_mir, Cat# 4427975, Applied Biosystems, Foster City, CA) TaqMan Advanced miRNA cDNA Synthesis Kit (Cat# A28007, Applied Biosystems, Foster City, CA) following the manufacturer’s protocol as described. RT-qPCR was performed using the TaqMan Fast Advanced Master Mix (Cat# 4444556, Applied Biosystems, Foster City, CA). U6 snRNA (Cat# 4444556, Applied Biosystems, Foster City, CA) was used for normalization.

RT-qPCR of mRNA was performed using the Applied Biosystems predesigned TaqMan Gene Expression Assays (Cat# 4331182, Applied Biosystems, Foster City, CA), and Superscript IV VILO Master Mix (Cat# 11766050, Invitrogen, Carlsbad, CA) per the manufacturer’s instructions for rat glyceraldehyde-3-phosphate dehydrogenase (Gapdh; Rn01462662_g1), transforming growth factor, beta 1 (Tgfb1; Rn00665219_g1), transforming growth factor, beta 2 (Tgfb1; Rn00579674_m1), zinc finger E-box binding homeobox 1 (Zeb1; Rn01538408_m1), zinc finger E-box binding homeobox 2 (Zeb2; Rn01449758_m1), SMAD family member 2 (Smad2; Rn01527104_g1) and SMAD family member 3 (Smad3; Rn01422011_m1).

### Lung Morphometry

Fetal rats were delivered by cesarean section on E21 and lung: body weight ratio was measured. For morphometric analysis, fetal lungs were fixed by intratracheal instillation of 4% paraformaldehyde under a constant pressure of 20 cm H_2_O. Visual confirmation of inflation was obtained for all specimens analyzed. After ligation of the trachea, the lungs were immersed in fixative and embedded in paraffin. Sections were stained with hematoxylin–eosin (H&E) or elastin van Gieson’s (EvG) stain. The slides were imaged on a Zeiss Axio Scope light microscope (Carl Zeiss Microscopy; Germany). Tissues were embedded in paraffin, and sliced sections from medial, mid- and lateral lung zones were mounted onto slides for H&E or EvG staining.^50^

Airspace enlargement was measured by assessing mean linear intercept (Lm). Three random fields were evaluated by microscopic projection and the alveolar size was estimated from the Lm of the airspace as described previously.^51–54^ Alveolar thickness was measured as the mean septal wall thickness of 10 terminal alveoli per field.

EvG stained sections were used for pulmonary arteriole (PA) remodeling assessment. The PAs were distinguished from pulmonary veins based on their position and structure. In each sample, only small PAs (external diameter: 20 to 100 μm) were used. External diameter (ED) was measured across the shortest luminal profile between the external elastic laminae. Diameter across the longest luminal profile also was measured, and only arteries in which the longest diameter did not exceed ED by more than 50% were used. Medial thickness was measured on the same profile as ED. To assess PA remodeling, the percentage of the medial wall thickness was calculated according to the following formula: (2 × medial wall thickness/external diameter) × 100% as previously described.^55–57^

### Statistical Analysis

Data are presented as means ± S.D. unless otherwise noted and compared using student’s t-test, or one-way or two-way ANOVA followed by Tukey’s post-hoc test for multiple comparisons when appropriate. Bonferroni correction was used to correct for multiple comparisons. Statistical analyses were carried out using GraphPad Prism v9. A *P* value of less than 0.05 was considered statistically significant.

## Acknowledgements

This work was supported by the National Institutes of Health (UG3-HL147352 to W.M.S., K08HL135402 to M.S.). A.S.P. was supported by two NIH National Research Service Awards (NRSAs) (a T32 GM86287 training grant and a F32 HL142144 individual postdoctoral fellowship), as well as a postdoctoral research fellowship award (PIOTRO20F0) from the Cystic Fibrosis Foundation (CFF). S.J.U. was supported by a Cystic Fibrosis Foundation award (STITEL17G0). S.L.A was supported by a Yale Department of Surgery Ohse Award.

Authors S.J.U., W.M.S., A.S.R., D.H.S., and A.S.P. have filed an invention disclosure for the technology used in this manuscript with the Yale Office of Cooperative Research.

Authors W.M.S., A.S.R., D.H.S., A.S.P. are listed as inventors on the following patent: https://patents.justia.com/patent/20200113821

The graphical abstract for this manuscript was created using Mind The Graph: https://mindthegraph.com/

## Author Contributions

S.J.U. – conceptualization, data curation, formal analysis, investigation, methodology, project administration, visualization, writing – original draft, writing – review & editing

N.K.Y. – investigation, project administration, visualization

T.J.P. – data curation, investigation

N.L.M. – investigation

M.E.G. – formal analysis

M.F.W. – investigation

S.L.A – investigation, funding acquisition

A.S.R. – conceptualization, methodology, supervision

M.S. – formal analysis, funding acquisition, methodology, supervision, writing – review & editing

W.M.S. – conceptualization, funding acquisition, methodology, resources, supervision, writing – review & editing

A.S.P. – conceptualization, funding acquisition, investigation, methodology, supervision, writing – review & editing

D.H.S. – conceptualization, funding acquisition, methodology, project administration, resources, supervision, writing – review & editing

## References

1 Longoni, M., High, F. A., Russell, M. K., Kashani, A., Tracy, A. A., Coletti, C. M., Hila, R., Shamia, A., Wells, J., Ackerman, K. G. et al. Molecular pathogenesis of congenital diaphragmatic hernia revealed by exome sequencing, developmental data, and bioinformatics. Proc Natl Acad Sci U S A 111, 12450–12455, doi:10.1073/pnas.1412509111 (2014).

2 Veenma, D. C., de Klein, A. & Tibboel, D. Developmental and genetic aspects of congenital diaphragmatic hernia. Pediatr Pulmonol 47, 534–545, doi:10.1002/ppul.22553 (2012).

3 Langham, M. R., Kays, D. W., Ledbetter, D. J., Frentzen, B., Sanford, L. L. & Richards, D. S. Congenital Diaphragmatic Hernia: Epidemiology and Outcome. Clinics in Perinatology 23, 671–688, doi:https://doi.org/10.1016/S0095-5108(18)30201-X (1996).

4 Wynn, J., Krishnan, U., Aspelund, G., Zhang, Y., Duong, J., Stolar, C. J., Hahn, E., Pietsch, J., Chung, D., Moore, D. et al. Outcomes of congenital diaphragmatic hernia in the modern era of management. J Pediatr 163, 114–119 e111, doi:10.1016/j.jpeds.2012.12.036 (2013).

5 Coughlin, M. A., Werner, N. L., Gajarski, R., Gadepalli, S., Hirschl, R., Barks, J., Treadwell, M. C., Ladino-Torres, M., Kreutzman, J. & Mychaliska, G. B. Prenatally diagnosed severe CDH: mortality and morbidity remain high. J Pediatr Surg 51, 1091–1095, doi:10.1016/j.jpedsurg.2015.10.082 (2016).

6 McGivern, M. R., Best, K. E., Rankin, J., Wellesley, D., Greenlees, R., Addor, M. C., Arriola, L., de Walle, H., Barisic, I., Beres, J. et al. Epidemiology of congenital diaphragmatic hernia in Europe: a register-based study. Arch Dis Child Fetal Neonatal Ed 100, F137–144, doi:10.1136/archdischild-2014-306174 (2015).

7 Ameis, D., Khoshgoo, N. & Keijzer, R. Abnormal lung development in congenital diaphragmatic hernia. Seminars in Pediatric Surgery 26, 123–128, doi:https://doi.org/10.1053/j.sempedsurg.2017.04.011 (2017).

8 Donahoe, P. K., Longoni, M. & High, F. A. Polygenic Causes of Congenital Diaphragmatic Hernia Produce Common Lung Pathologies. The American Journal of Pathology 186, 2532–2543, doi:https://doi.org/10.1016/j.ajpath.2016.07.006 (2016).

9 George, D. K., Cooney, T. P., Chiu, B. K. & Thurlbeck, W. M. Hypoplasia and Immaturity of the Terminal Lung Unit (Acinus) in Congenital Diaphragmatic Hernia. American Review of Respiratory Disease 136, 947–950, doi:10.1164/ajrccm/136.4.947 (1987).

10 Kool, H., Mous, D., Tibboel, D., de Klein, A. & Rottier, R. J. Pulmonary vascular development goes awry in congenital lung abnormalities. Birth Defects Research Part C: Embryo Today: Reviews 102, 343–358, doi:10.1002/bdrc.21085 (2014).

11 Deprest, J. A., Nicolaides, K. H., Benachi, A., Gratacos, E., Ryan, G., Persico, N., Sago, H., Johnson, A., Wielgoś, M., Berg, C. et al. Randomized Trial of Fetal Surgery for Severe Left Diaphragmatic Hernia. The New England journal of medicine, doi:10.1056/NEJMoa2027030 (2021).

12 Deprest, J. A., Benachi, A., Gratacos, E., Nicolaides, K. H., Berg, C., Persico, N., Belfort, M., Gardener, G. J., Ville, Y., Johnson, A. et al. Randomized Trial of Fetal Surgery for Moderate Left Diaphragmatic Hernia. The New England journal of medicine, doi:10.1056/NEJMoa2026983 (2021).

13 Pereira-Terra, P., Deprest, J. A., Kholdebarin, R., Khoshgoo, N., DeKoninck, P., Munck, A. A., Wang, J., Zhu, F., Rottier, R. J., Iwasiow, B. M. et al. Unique Tracheal Fluid MicroRNA Signature Predicts Response to FETO in Patients With Congenital Diaphragmatic Hernia. Ann Surg 262, 1130–1140, doi:10.1097/sla.0000000000001054 (2015).

14 Burk, U., Schubert, J., Wellner, U., Schmalhofer, O., Vincan, E., Spaderna, S. & Brabletz, T. A reciprocal repression between ZEB1 and members of the miR-200 family promotes EMT and invasion in cancer cells. EMBO Rep 9, 582–589, doi:10.1038/embor.2008.74 (2008).

15 Gregory, P. A., Bracken, C. P., Smith, E., Bert, A. G., Wright, J. A., Roslan, S., Morris, M., Wyatt, L., Farshid, G., Lim, Y. Y. et al. An autocrine TGF-beta/ZEB/miR-200 signaling network regulates establishment and maintenance of epithelial-mesenchymal transition. Mol Biol Cell 22, 1686–1698, doi:10.1091/mbc.E11-02-0103 (2011).

16 Lu, J., Qian, J., Izvolsky, K. I. & Cardoso, W. V. Global analysis of genes differentially expressed in branching and non-branching regions of the mouse embryonic lung. Developmental biology 273, 418–435, doi:10.1016/j.ydbio.2004.05.035 (2004).

17 Khoshgoo, N., Visser, R., Falk, L., Day, C. A., Ameis, D., Iwasiow, B. M., Zhu, F., Ozturk, A., Basu, S., Pind, M. et al. MicroRNA-200b regulates distal airway development by maintaining epithelial integrity. Sci Rep 7, 6382, doi:10.1038/s41598-017-05412-y (2017).

18 Khoshgoo, N., Kholdebarin, R., Pereira-Terra, P., Mahood, T. H., Falk, L., Day, C. A., Iwasiow, B. M., Zhu, F., Mulhall, D., Fraser, C. et al. Prenatal microRNA miR-200b Therapy Improves Nitrofen-induced Pulmonary Hypoplasia Associated With Congenital Diaphragmatic Hernia. Annals of surgery 269, 979–987, doi:10.1097/sla.0000000000002595 (2019).

19 Rupaimoole, R. & Slack, F. J. MicroRNA therapeutics: towards a new era for the management of cancer and other diseases. Nat Rev Drug Discov 16, 203–222, doi:10.1038/nrd.2016.246 (2017).

20 Rupaimoole, R., Han, H. D., Lopez-Berestein, G. & Sood, A. K. MicroRNA therapeutics: principles, expectations, and challenges. Chin J Cancer 30, 368–370, doi:10.5732/cjc.011.10186 (2011).

21 Li, Z. & Rana, T. M. Therapeutic targeting of microRNAs: current status and future challenges. Nat Rev Drug Discov 13, 622–638, doi:10.1038/nrd4359 (2014).

22 van Rooij, E. & Kauppinen, S. Development of microRNA therapeutics is coming of age. EMBO Mol Med 6, 851–864, doi:10.15252/emmm.201100899 (2014).

23 Blum, J. S. & Saltzman, W. M. High loading efficiency and tunable release of plasmid DNA encapsulated in submicron particles fabricated from PLGA conjugated with poly-L-lysine. J Control Release 129, 66–72, doi:10.1016/j.jconrel.2008.04.002 (2008).

24 Trang, P., Wiggins, J. F., Daige, C. L., Cho, C., Omotola, M., Brown, D., Weidhaas, J. B., Bader, A. G. & Slack, F. J. Systemic delivery of tumor suppressor microRNA mimics using a neutral lipid emulsion inhibits lung tumors in mice. Mol Ther 19, 1116–1122, doi:10.1038/mt.2011.48 (2011).

25 Ozpolat, B., Sood, A. K. & Lopez-Berestein, G. Nanomedicine based approaches for the delivery of siRNA in cancer. J Intern Med 267, 44–53, doi:10.1111/j.1365-2796.2009.02191.x (2010).

26 Joshi, H. P., Subramanian, I. V., Schnettler, E. K., Ghosh, G., Rupaimoole, R., Evans, C., Saluja, M., Jing, Y., Cristina, I., Roy, S. et al. Dynamin 2 along with microRNA-199a reciprocally regulate hypoxia-inducible factors and ovarian cancer metastasis. Proc Natl Acad Sci USA 111, 5331–5336, doi:10.1073/pnas.1317242111 (2014).

27 Dahlman, J. E., Barnes, C., Khan, O. F., Thiriot, A., Jhunjunwala, S., Shaw, T. E., Xing, Y., Sager, H. B., Sahay, G., Speciner, L. et al. In vivo endothelial siRNA delivery using polymeric nanoparticles with low molecular weight. Nature Nanotechnology 9, 648–655, doi:10.1038/nnano.2014.84 (2014).

28 Woodrow, K. A., Cu, Y., Booth, C. J., Saucier-Sawyer, J. K., Wood, M. J. & Mark Saltzman, W. Intravaginal gene silencing using biodegradable polymer nanoparticles densely loaded with small-interfering RNA. Nature Materials 8, 526–533, doi:10.1038/nmat2444 (2009).

29 Piotrowski-Daspit, A. S., Kauffman, A. C., Bracaglia, L. G. & Saltzman, W. M. Polymeric vehicles for nucleic acid delivery. Advanced drug delivery reviews 156, 119–132, doi:10.1016/j.addr.2020.06.014 (2020).

30 Lv, H., Zhang, S., Wang, B., Cui, S. & Yan, J. Toxicity of cationic lipids and cationic polymers in gene delivery. Journal of Controlled Release 114, 100–109, doi:https://doi.org/10.1016/j.jconrel.2006.04.014 (2006).

31 Makadia, H. K. & Siegel, S. J. Poly Lactic-co-Glycolic Acid (PLGA) as Biodegradable Controlled Drug Delivery Carrier. Polymers (Basel) 3, 1377–1397, doi:10.3390/polym3031377 (2011).

32 Zhou, J., Liu, J., Cheng, C. J., Patel, T. R., Weller, C. E., Piepmeier, J. M., Jiang, Z. & Saltzman, W. M. Biodegradable poly(amine-co-ester) terpolymers for targeted gene delivery. Nature Materials 11, 82–90, doi:10.1038/nmat3187 (2012).

33 Ullrich, S. J., Freedman-Weiss, M., Ahle, S., Mandl, H. K., Piotrowski-Daspit, A. S., Roberts, K., Yung, N., Maassel, N., Bauer-Pisani, T., Ricciardi, A. S. et al. Nanoparticles for delivery of agents to fetal lungs. Acta biomaterialia 123, 346–353, doi:10.1016/j.actbio.2021.01.024 (2021).

34 Ricciardi, A. S., Bahal, R., Farrelly, J. S., Quijano, E., Bianchi, A. H., Luks, V. L., Putman, R., Lopez-Giraldez, F., Coskun, S., Song, E. et al. In utero nanoparticle delivery for site-specific genome editing. Nat Commun 9, 2481, doi:10.1038/s41467-018-04894-2 (2018).

35 Schittny, J. C. Development of the lung. Cell Tissue Res 367, 427–444, doi:10.1007/s00441-016-2545-0 (2017).

36 Alejandre-Alcázar, M. A., Michiels-Corsten, M., Vicencio, A. G., Reiss, I., Ryu, J., de Krijger, R. R., Haddad, G. G., Tibboel, D., Seeger, W., Eickelberg, O. et al. TGF-beta signaling is dynamically regulated during the alveolarization of rodent and human lungs. Developmental dynamics: an official publication of the American Association of Anatomists 237, 259–269, doi:10.1002/dvdy.21403 (2008).

37 Chen, H., Sun, J., Buckley, S., Chen, C., Warburton, D., Wang, X. F. & Shi, W. Abnormal mouse lung alveolarization caused by Smad3 deficiency is a developmental antecedent of centrilobular emphysema. American journal of physiology. Lung cellular and molecular physiology 288, L683–691, doi:10.1152/ajplung.00298.2004 (2005).

38 Mous, D. S., Buscop-van Kempen, M. J., Wijnen, R. M. H., Tibboel, D., Morty, R. E. & Rottier, R. J. Opposing Effects of TGFβ and BMP in the Pulmonary Vasculature in Congenital Diaphragmatic Hernia. Frontiers in medicine 8, 642577, doi:10.3389/fmed.2021.642577 (2021).

39 Kauffman, A. C., Piotrowski-Daspit, A. S., Nakazawa, K. H., Jiang, Y., Datye, A. & Saltzman, W. M. Tunability of Biodegradable Poly(amine-co-ester) Polymers for Customized Nucleic Acid Delivery and Other Biomedical Applications. Biomacromolecules 19, 3861–3873, doi:10.1021/acs.biomac.8b00997 (2018).

40 Chinoy, M. R. Lung growth and development. Frontiers in bioscience: a journal and virtual library 8, d392–415, doi:10.2741/974 (2003).

41 Bates, S. E. Epigenetic Therapies for Cancer. The New England journal of medicine 383, 650–663, doi:10.1056/NEJMra1805035 (2020).

42 Wagner, R., Montalva, L., Zani, A. & Keijzer, R. Basic and translational science advances in congenital diaphragmatic hernia. Seminars in perinatology 44, 151170, doi:10.1053/j.semperi.2019.07.009 (2020).

43 Van Kamp, I. L., Klumper, F. J., Oepkes, D., Meerman, R. H., Scherjon, S. A., Vandenbussche, F. P. & Kanhai, H. H. Complications of intrauterine intravascular transfusion for fetal anemia due to maternal red-cell alloimmunization. American journal of obstetrics and gynecology 192, 171–177, doi:10.1016/j.ajog.2004.06.063 (2005).

44 Bang, J., Bock, J. E. & Trolle, D. Ultrasound-guided fetal intravenous transfusion for severe rhesus haemolytic disease. Br Med J (Clin Res Ed) 284, 373–374 (1982).

45 Pasman, S. A., Claes, L., Lewi, L., Van Schoubroeck, D., Debeer, A., Emonds, M., Geuten, E., De Catte, L. & Devlieger, R. Intrauterine transfusion for fetal anemia due to red blood cell alloimmunization: 14 years experience in Leuven. Facts, views & vision in ObGyn 7, 129–136 (2015).

46 Perrone, E. E. & Deprest, J. A. Fetal endoscopic tracheal occlusion for congenital diaphragmatic hernia: a narrative review of the history, current practice, and future directions. Translational pediatrics 10, 1448–1460, doi:10.21037/tp-20-130 (2021).

47 Grun, M. K., Suberi, A., Shin, K., Lee, T., Gomerdinger, V., Moscato, Z. M., Piotrowski-Daspit, A. S. & Saltzman, W. M. PEGylation of poly(amine-co-ester) polyplexes for tunable gene delivery. Biomaterials 272, 120780, doi:10.1016/j.biomaterials.2021.120780 (2021).

48 Cui, J., Piotrowski-Daspit, A. S., Zhang, J., Shao, M., Bracaglia, L. G., Utsumi, T., Seo, Y. E., DiRito, J., Song, E., Wu, C. et al. Poly(amine-co-ester) nanoparticles for effective Nogo-B knockdown in the liver. J Control Release 304, 259–267, doi:10.1016/j.jconrel.2019.04.044 (2019).

49 Kluth, D., Kangah, R., Reich, P., Tenbrinck, R., Tibboel, D. & Lambrecht, W. Nitrofen-induced diaphragmatic hernias in rats: an animal model. J Pediatr Surg 25, 850–854, doi:10.1016/0022-3468(90)90190-k (1990).

50 Burgos, C. M., Pearson, E. G., Davey, M., Riley, J., Jia, H., Laje, P., Flake, A. W. & Peranteau, W. H. Improved pulmonary function in the nitrofen model of congenital diaphragmatic hernia following prenatal maternal dexamethasone and/or sildenafil. Pediatric research 80, 577–585, doi:10.1038/pr.2016.127 (2016).

51 Knudsen, L., Weibel, E. R., Gundersen, H. J., Weinstein, F. V. & Ochs, M. Assessment of air space size characteristics by intercept (chord) measurement: an accurate and efficient stereological approach. J Appl Physiol (1985) 108, 412–421, doi:10.1152/japplphysiol.01100.2009 (2010).

52 Thurlbeck, W. M. Internal surface area and other measurements in emphysema. Thorax 22, 483–496, doi:10.1136/thx.22.6.483 (1967).

53 Kim, S. J., Shan, P., Hwangbo, C., Zhang, Y., Min, J. N., Zhang, X., Ardito, T., Li, A., Peng, T., Sauler, M. et al. Endothelial toll-like receptor 4 maintains lung integrity via epigenetic suppression of p16(INK4a). Aging Cell 18, e12914, doi:10.1111/acel.12914 (2019).

54 Hsia, C. C., Hyde, D. M., Ochs, M. & Weibel, E. R. An official research policy statement of the American Thoracic Society/European Respiratory Society: standards for quantitative assessment of lung structure. American journal of respiratory and critical care medicine 181, 394–418, doi:10.1164/rccm.200809-1522ST (2010).

55 Kanai, M., Kitano, Y., von Allmen, D., Davies, P., Adzick, N. S. & Flake, A. W. Fetal tracheal occlusion in the rat model of nitrofen-induced congenital diaphragmatic hernia: tracheal occlusion reverses the arterial structural abnormality. J Pediatr Surg 36, 839–845, doi:10.1053/jpsu.2001.23950 (2001).

56 Umeda, S., Miyagawa, S., Fukushima, S., Oda, N., Saito, A., Sakai, Y., Sawa, Y. & Okuyama, H. Enhanced Pulmonary Vascular and Alveolar Development via Prenatal Administration of a Slow-Release Synthetic Prostacyclin Agonist in Rat Fetal Lung Hypoplasia. PLoS One 11, e0161334–e0161334, doi:10.1371/journal.pone.0161334 (2016).

57 Xu, X. F., Gu, W. Z., Wu, X. L., Li, R. Y. & Du, L. Z. Fetal pulmonary vascular remodeling in a rat model induced by hypoxia and indomethacin. J Matern Fetal Neonatal Med 24, 172–182, doi:10.3109/14767058.2010.482608 (2011).

